# Predicting primary production in the southern California Current Ecosystem from chlorophyll, nutrient concentrations, and irradiance

**DOI:** 10.1101/590240

**Authors:** Michael R. Stukel, Ralf Goericke, Michael R. Landry

**Affiliations:** Florida State University; University of California, San Diego

## Abstract

We investigated the processes driving variability in primary productivity in the California Current Ecosystem (CCE) in order to develop an algorithm for predicting primary productivity from in situ irradiance, nutrient, and chlorophyll (chl) measurements. Primary productivity data from seven process cruises of the CCE Long-Term Ecological Research (CCE LTER) program were used to parameterize the algorithm. An initial algorithm was developed using only irradiance to predict chl-specific productivity was found to have model-data misfit that was correlated with NH_4_^+^ concentrations. We thus found that the best estimates of primary productivity were obtained using an equation including NH_4_^+^ and irradiance: PP/Chl = V_0m_×(1-exp(−α×PAR/V_0m_)×NH_4_/(NH_4_+K_S_), where PP/Chl is chlorophyll-specific primary production in units of mg C d^−1^ / mg Chl, PAR is photosynthetically active radiation (units of μEi m^−2^ s^−1^), NH4+ is in units of μmol L^−1^, V_0m_ = 66.5 mg C d^−1^ / mg Chl, α = 1.5, and K_S_ = 0.025 μmol L^−1^. We then used this algorithm to compute primary productivity rates for the CCE-P1706 cruise on which in situ primary productivity samples were not available. We compared these estimates to independent productivity estimates derived from protistan grazing dilution experiments and found excellent agreement.

## I – Algorithm Development

Morrow et al. (2018) tested the light dependency of phytoplankton specific growth rates by using ^14^C-PP divided by Chl as a proxy for phytoplankton specific growth rates. They used an equation of the form:

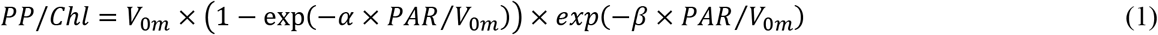

Because they were specifically focused on the ecosystem responses to interannual variability (especially El Niño), they restricted their study to measurements from the P0605, P0704, P0810, P1408, and P1604 cruises and found best fit parameters of: V_0m_ = 64 mg C d^−1^ / mg Chl, α = 1.0, and β = 0.049, when light was given as daily average PAR values in units of μEi m^−2^ s^−1^. We updated these analyses by including additional data from the P1106 and P1208 “front” cruises (Krause et al. 2015; Stukel et al. 2017). With this additional data, the best fit parameter values were: V_0m_ = 52 mg C d^−1^ / mg Chl, α = 1.2, and β = 0.011. We then investigated the relationship between model-data misfit and nutrient concentrations to investigate whether or not nutrient limitation played an additional role in determining phytoplankton specific growth rates (Fig. 2). Although no statistically significant relationship was found between percent error and either nitrate concentration or total dissolved inorganic nitrogen (NO_3_ + NO_2_ + NH_4_), a relationship was found between percent error and ammonium concentration (non-parametric Spearman’s ρ = 0.31, p = 1.3×10^12^). This suggests that Eq. 1 overestimates primary productivity when ammonium concentrations are low and underestimates it when ammonium concentrations are high.

**Fig. 1.**
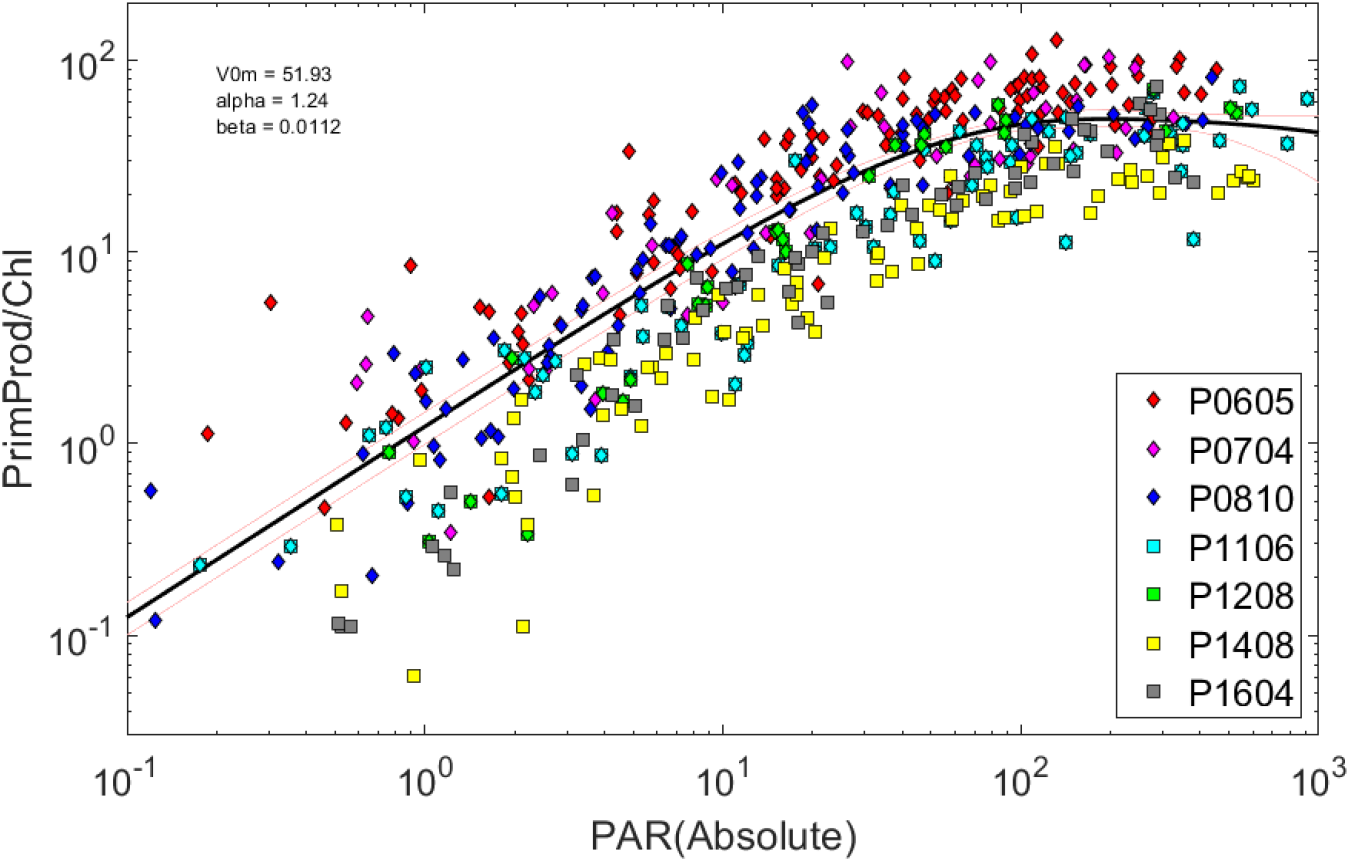
Relationship between PAR (μEi m^−2^ s^−1^) and the ratio of ^14^C-PP to Chl a (mg C d^−1^ / mg Chl) as determined on 6 CCE LTER Process cruises.

**Fig. 2.**
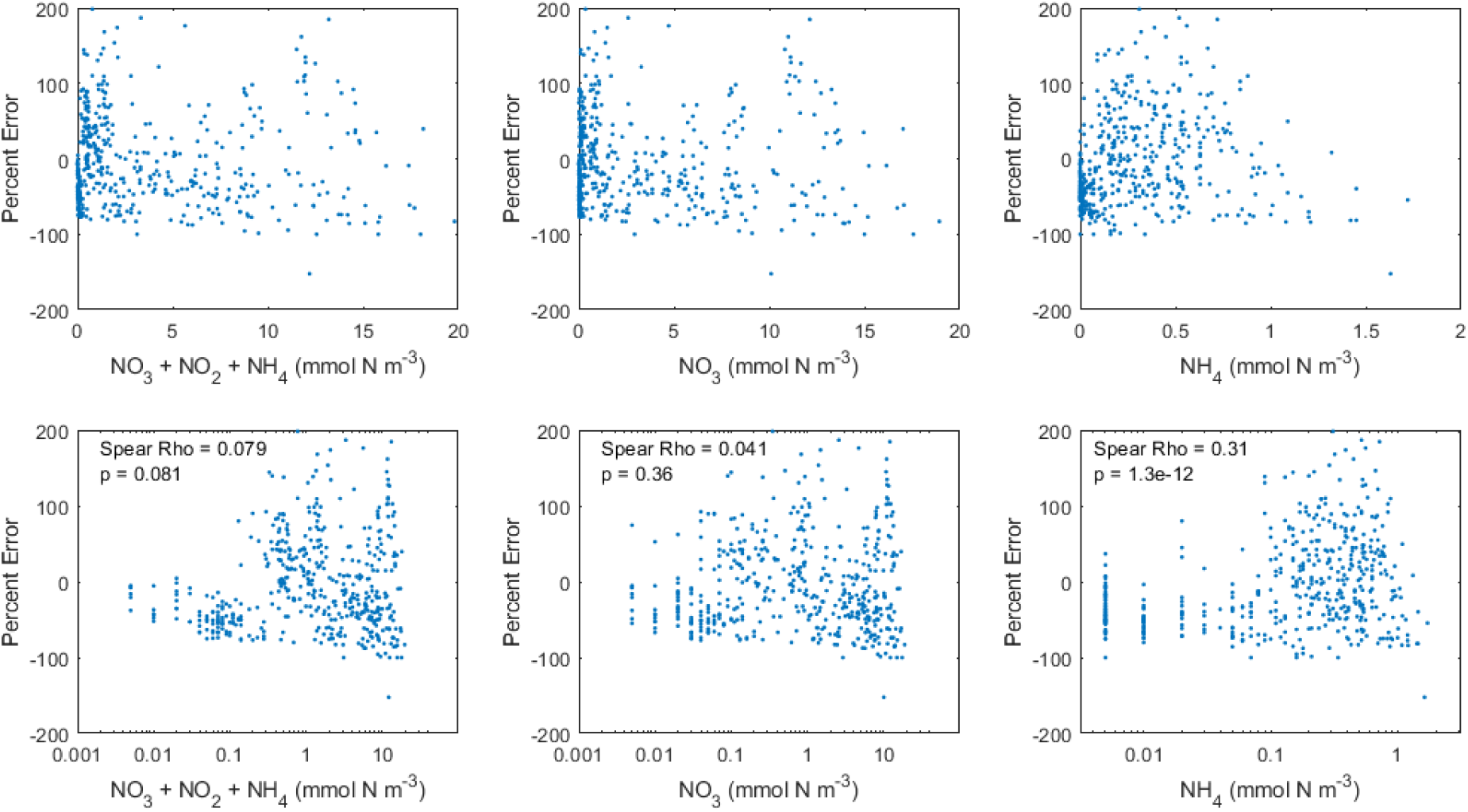
Model data misfits plotted against nutrient concentration. Bottom plots are logarithmic scaling of nutrients.

We chose to incorporate ammonium limitation of primary productivity by modeling it as a Monod function of the form: LIM_NH4_ = NH_4_/(NH_4_+K_S_). Because this function will, by definition, set primary productivity equal to zero when NH_4_^+^ values are reported as 0 μmol L^−1^ (i.e., below the detection limit), we assumed that when NH_4_^+^ was reported as 0 μmol L^−1^, its actual in situ value was equal to one half of the detection limit (in other words, we replaced all 0 values as 5 nmol L^−1^). When all of the data from P0605, P0704, P0810, P1106, P1208, P1408, and P1604 was fit to a model of the form:

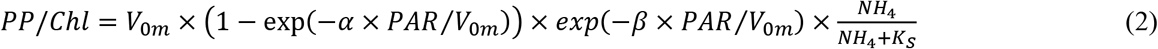

(with the constraints that K_S_, V_0m_, α, and β were all ≥0) we obtained values of V_0m_ = 66.5 mg C d^−1^ / mg Chl, α = 1.5, β = 0.0, and K_S_ = 0.025. We thus concluded that the photoinhibition term did not need to be included if NH_4_^+^ was used as a predictor variable and further tested the equation:

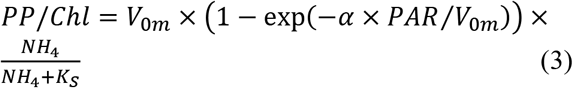

To determine whether or not the inclusion of NH_4_^+^ as a predictor variable actually increased the predictive ability of the model, we conducted a Monte Carlo analysis in which 50% of the data points were selected at random (a training data set) and used to parameterize both Eq. 1 and Eq. 3. Model data misfits were then computed with respect to how well each algorithm predicted the half of the data points that had been withheld from the training dataset. This process was repeated 1000 times and showed that the inclusion of NH_4_^+^ as a predictor variable typically reduced the sum of squared misfits by ~25%. The model with NH_4_^+^ was a better predictor with all of the training sets.

We then quantified the uncertainty in primary productivity predicted by the algorithm by computing model-data misfits for all data points. We found (unsurprisingly) that absolute misfit was strongly correlated with both the predicted and measured value of primary productivity. However, the relative misfit (PercentError = (measurement − estimate)/estimate×100%) was not correlated with measured primary productivity. Hence, we can use a constant PercentError value to determine uncertainty limits on our estimates. PercentError was not symmetric with respect to upper and lower limits. Rather, we found that 95% of measured values fell between 14% and 262% of the predicted value (Fig. 4). Similarly, 68% of measured values (i.e., one standard deviation) fell between 44% and 161% of the predicted value). We also computed the correlation between PercentError values for primary productivity measurements made at multiple depths on the same CTD casts. The median correlation across all casts was 0.18. For measurements made at the same depths on different casts within the same cycle, the correlation in PercentError was 0.15. These correlation values were included in error propagation when determining vertically-integrated primary productivity or cycle average productivity at specific depths.

**Fig. 3.**
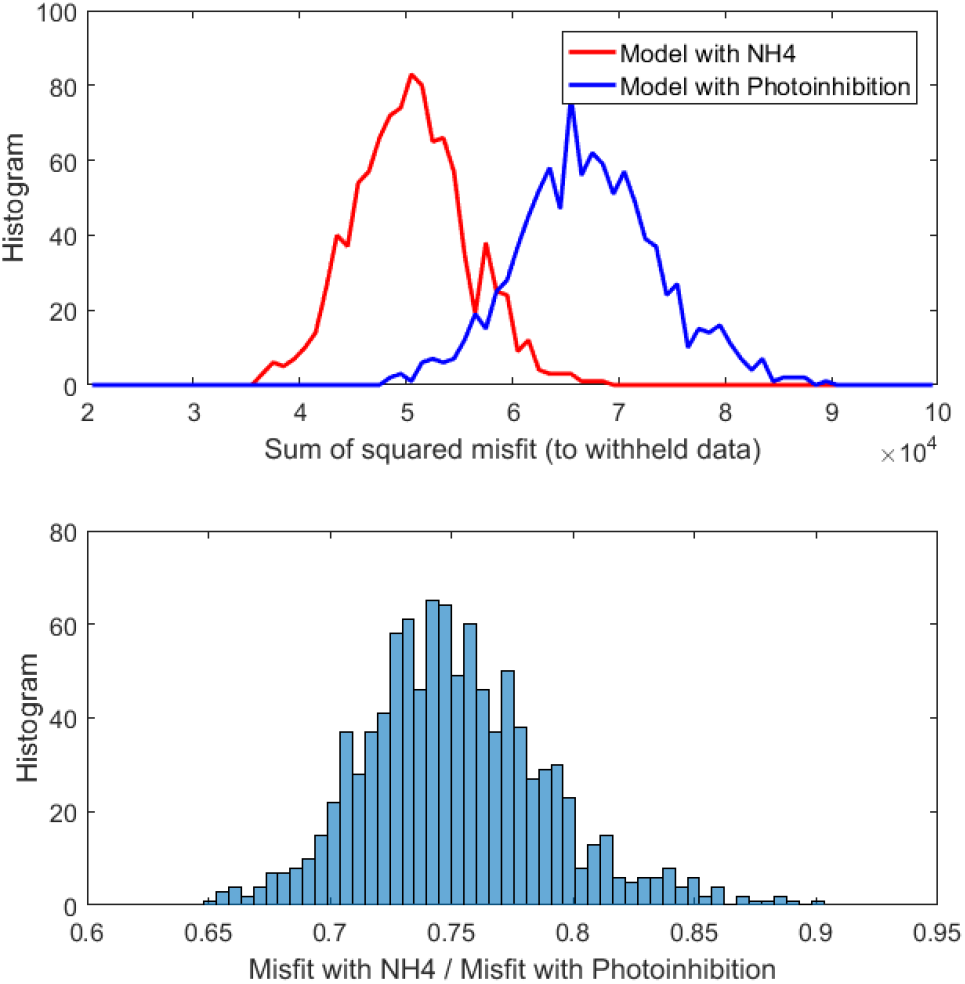
Model data misfit for Monte Carlo simulations using 50% of data as a training set and testing against the withheld data points. Model with NH_4_ uses Eq. 3. Model with Photoinhibition uses Eq. 1.

**Fig. 4.**
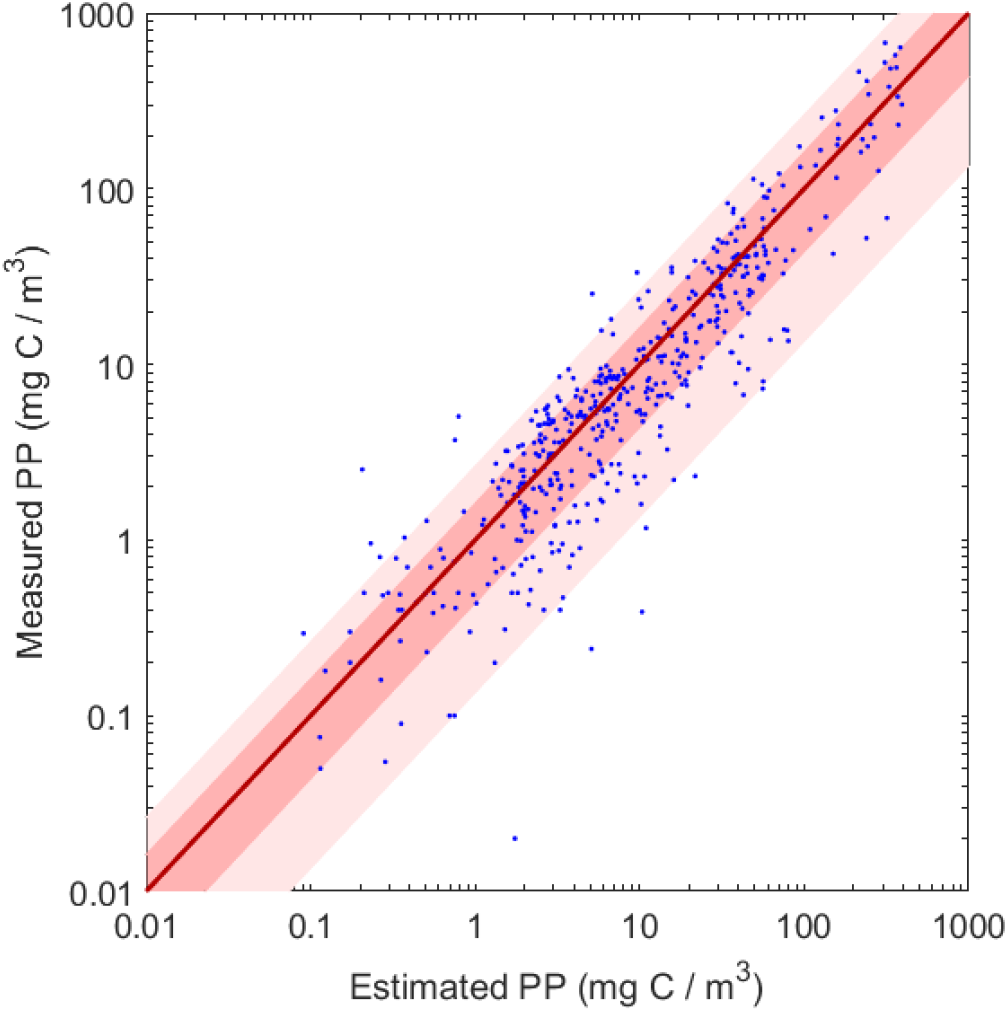
Comparison between model estimated primary productivity (Eq. 3) and measured primary productivity. Thick red line is the 1:1 line. Dark pink shows 68% confidence limits. Light pink shows 95% confidence limits.

## II – Application to P1706 Field Data

Chlorophyll concentrations were measured by R. Goericke using the acidification method with samples taken either from Niskin bottles (cycle measurements and measurements made on cross-feature or along-feature transects) or from the ship’s flow-through system (surface samples taken during SeaSoar surveys). Nutrient concentrations were measured by autoanalyzer on 50-mL frozen samples that had been filtered at sea through a 0.1-μm cartridge filter. Surface PAR was determined as daily average for the 24-hours following sample collection (to closely match assumptions used when conducting bottle incubations on previous cruises). PAR data was available from the R.V. Revelle’s meteorological (MET) system. For cycle data, percent surface irradiance (PSI) was determined as the average PSI at sample collection depth on daytime CTD casts collected during the 24-hours following sample collection. CTD data was retrieved from the CCE LTER DataZoo website and had been processed by R. Goericke. During transect sampling PSI data was typically not available, because we did not spend 24 hours at a particular station. Thus for transect data we computed PSI using beam transmission data. For casts on which beam transmission and ambient PAR were available, we computed the light extinction coefficient and regressed the light extinction coefficient against beam transmission (R^2^ = 0.33, p ≪ 10^−10^). This relationship between light extinction coefficient and beam transmission was then used to compute depth-varying light extinction coefficients and PSI for casts when PSI was not directly measured by CTD. In a comparison of primary productivity computed using the beam-c data to primary productivity computed using the in situ PAR measurements, the R^2^ was 0.88 with p ≪ 10^−10^. We then applied Eq. 3 and multiplied by Chl to determine primary productivity estimates (Fig. 5). Error in Eq. 3 was propagated to individual data points and summary data.

**Fig. 5.**
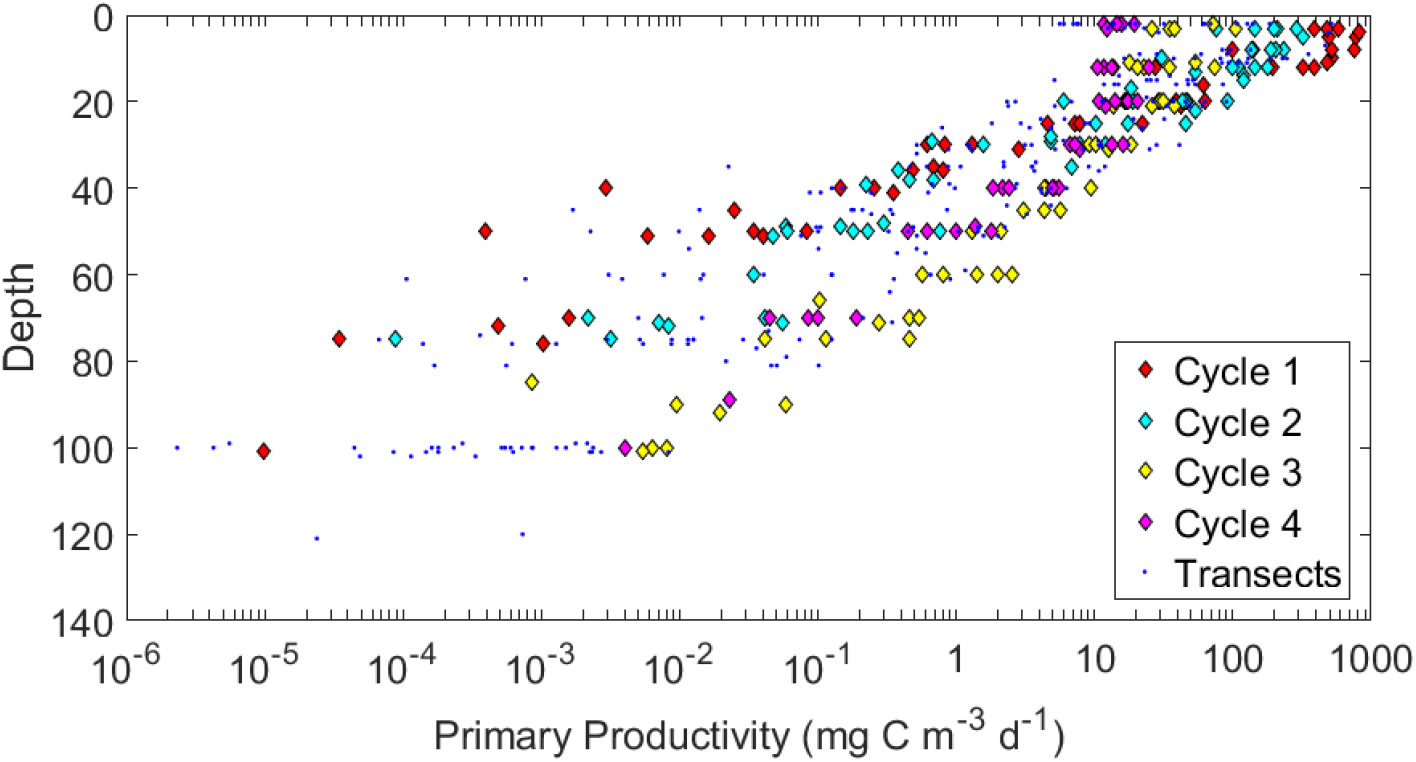
Primary productivity as a function of depth on the P1706 Cruise, computed using Eq. 3.

## III – Model validation with independent growth rate measurements

Phytoplankton growth rate estimates were measured daily at six depths *in situ* during Lagrangian cycles. Growth rate measurements were made using two-point “mini-dilution” protistan grazing experiments (Landry et al. 1984; Landry et al. 2008). Experiments were conducted in 2.7-L polycarbonate bottles placed inside mesh bags hanging off of an *in situ* array with a 3×1-m drogue centered in the mixed layer (Landry et al. 2009; Landry et al. 2012). These experiments provide chlorophyll-based growth rate estimates. To convert these estimates to carbon-based productivity, we need to know the Chl a concentration (measured during the experiment) and the C:Chl ratio of the ambient phytoplankton community, which was not determined experimentally. To estimate C:Chl ratios, we used the Geider et al. (1997) model as modified and parameterized for the CCE by Li et al. (2010). We first used this algorithm to predict C:Chl ratios for the P0605, P0704 and P0810 cruises, because direct microscopy-based estimates of C:Chl *in situ* C:Chl ratios were available for these cruises (Taylor et al. 2012). We found that at the 95% confidence limit, *in situ* measurements were between 25% and 462% of the estimated value, while at the 68% confidence interval (i.e., one standard deviation) the *in situ* measurements were between 44% and 178% of the estimated value.

Comparison between the algorithm-based and dilution-based estimates of primary productivity showed excellent agreement (Fig. 6). When only comparing paired samples, the mean primary productivity estimated using the algorithm was 120 mg C m^−3^ d^−1^ with a standard deviation of 190 mg C m^−3^ d^−1^; for the dilution-based estimates the mean was 111 mg C m^−3^ d^−1^ with a standard deviation of 206 mg C m^−3^ d^−1^. The non-parametric Spearman’s rank correlation between the two estimates was 0.86 (p≪10^−6^). The median misfit between the two estimates was 1.9 mg C m^−2^ d^−1^, suggesting negligible biases between the two approaches. The median absolute value of the misfit was 9.6, mg C m^−2^ d^−1^, while the median absolute value of the percent error was 41%.

**Fig. 6.**
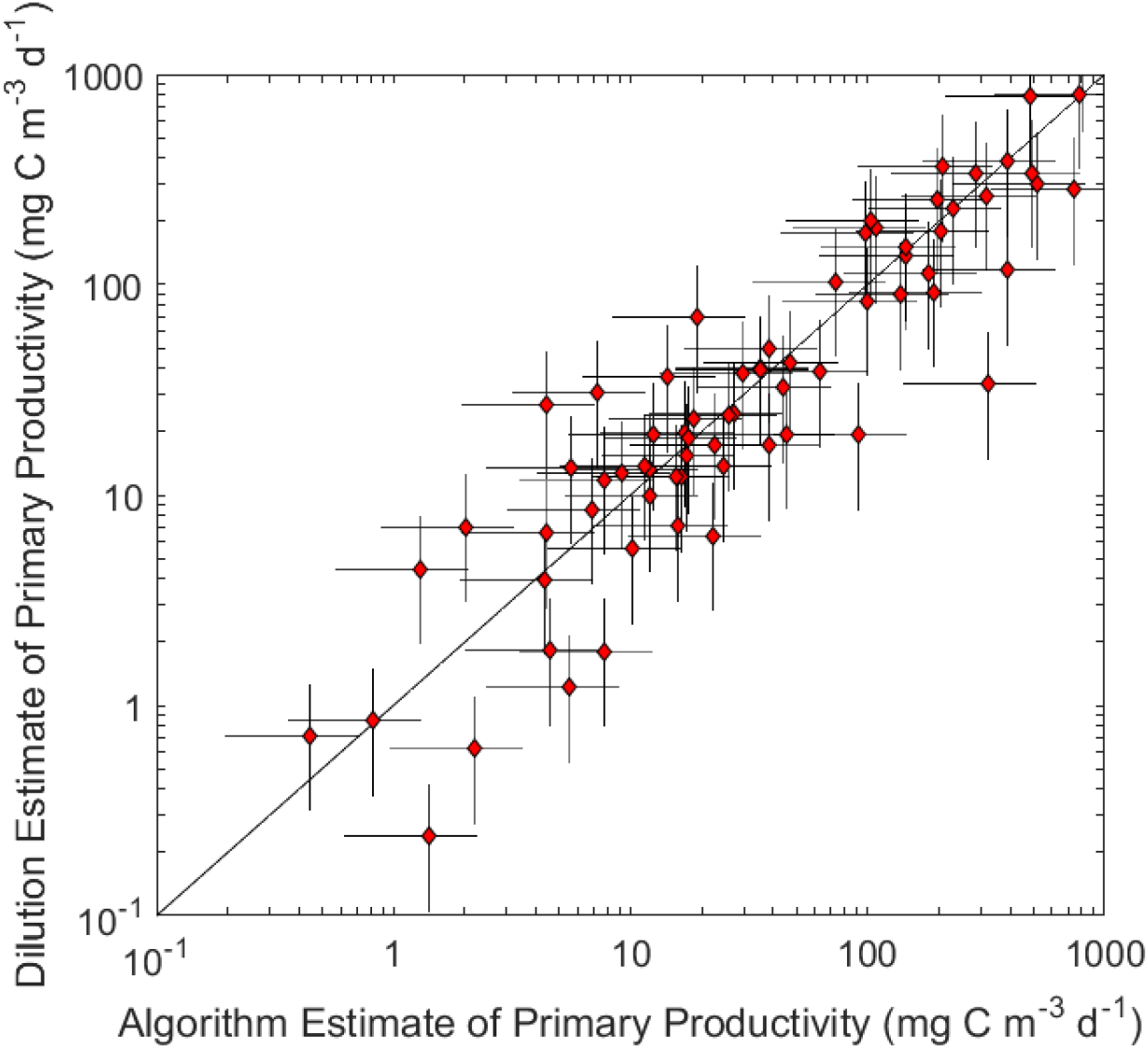
Comparison of CCE-P1706 primary productivity estimated with our algorithm (Eq. 3, x-axis) and primary productivity estimated from protistan grazing dilution experiments (y-axis). Error bars equate to one standard deviation of the uncertainty in each measurement.

## ACKNOWLEDGMENTS

We thank the captains and crews of the R.V. Melville, R. V. Revelle, R. V. Knorr, R. V. Thompson, and R. V. Sikuliaq as well as our many collaborators in the CCE LTER Program. Data used in this manuscript can be found on the CCE LTER DataZoo website: https://oceaninformatics.ucsd.edu/datazoo/catalogs/ccelter/datasets. This work was funded by NSF Bio Oce grants to the CCE LTER Program: OCE-0417616, OCE-1026607, OCE-1637632, and OCE-1614359.

